# SynBioGPT: A Retrieval-Augmented Large Language Model Platform for AI-Guided Microbial Strain Development

**DOI:** 10.1101/2025.03.23.644789

**Authors:** Zhitao Mao, Jun Du, Ruoyu Wang, Haoran Li, Jirun Guan, Zhenkun Shi, Xiaoping Liao, Hongwu Ma

**Author notes:** These authors contributed equally to this work (Zhitao Mao, Jun Du, Ruoyu Wang, Haoran Li).

## Abstract

Synthetic biology seeks to engineer microbial cell factories for sustainable bioproduction, yet the optimization of these systems is impeded by the complexity of metabolic engineering and the protracted timelines of iterative design-build-test-learn (DBTL) cycles. Traditional computational approaches, such as constraint-based modeling, provide valuable insights but demand extensive manual curation. Large language models (LLMs) hold promises for automating knowledge extraction and strain design, yet conventional models like GPT-4 suffer from outdated corpora and hallucination errors in domain-specific tasks. SynBioGPT v1.0, a Retrieval-Augmented Generation (RAG)-enhanced LLM, enhanced knowledge retrieval using vector search but often retrieved semantically similar yet contextually irrelevant documents. Here, we introduce SynBioGPT v2.0 (https://synbiogpt.biodesign.ac.cn), which mitigates these limitations by decomposing queries into sub-questions and employing keyword-based searches. Tested on a 100-question synthetic biology benchmark, SynBioGPT v2.0 achieved 98% accuracy with the Claude-3.7-sonnet backend, a 10% improvement over v1.0’s 88% with Llama3-8B-Instruct. This advance highlight the efficacy of query decomposition and precise retrieval in enhancing LLM utility for synthetic biology.

## Introduction

Synthetic biology is an interdisciplinary field that applies engineering principles to design, construct, and control biological systems, enabling the predictive construction of programmable microbial cell factories for sustainable bioproduction and smart therapeutics [1]. Recent advances in automation, machine learning, and DNA synthesis have accelerated its applications in next-generation biorefineries [2]. A 2023 analysis highlights the increasing prioritization of metabolic engineering in synthetic biology research, with a growing focus on the biosynthesis of high-value compounds such as terpenoid-derived pharmaceuticals and biodegradable polymer precursors [3]. However, the industrial translation of these technologies is hindered by the time-intensive process of multi-parameter optimization of chassis strains, requiring extensive iterative design-build-test-learn (DBTL) cycles [4]. The complexity of metabolic engineering arises from the vast combinatorial space of genetic modifications, metabolic flux constraints, and regulatory interactions, making it challenging to predict optimal engineering strategies [5]. Traditional computational approaches, including constraint-based modeling and kinetic simulations, offer insights but often require extensive manual curation and experimental validation, further slowing the engineering process [6].

To address these challenges, AI-driven tools have been increasingly integrated into synthetic biology workflows, aiming to automate knowledge extraction, optimize strain design, and enhance predictive capabilities [7, 8]. Machine learning models have shown promise in tasks such as pathway prediction, enzyme optimization, and regulatory network inference [9, 10]. However, conventional large language models (LLMs) such as GPT-4 [11] and LLaMA-3 [12] face significant challenges when applied to synthetic biology. First, these models rely on outdated or non-specialized training corpora [13], limiting their grasp of emerging metabolic pathways and novel bioprocesses. Second, they are prone to “hallucination” errors in the design of complex biochemical networks, potentially leading to inaccurate pathway predictions [14]. Retrieval-Augmented Generation (RAG) technology offers a compelling solution by combining a dynamically updated knowledge base with efficient data retrieval mechanisms, thereby ensuring precise and timely incorporation of domain-specific information [15]. We developed SynBioGPT v1.0, a RAG-enhanced LLM tailored for synthetic biology, leveraging a knowledge base of over 51,777 peer-reviewed studies [16]. However, its reliance on vector search—converting text into numerical vectors for similarity matching—can retrieve semantically similar but contextually irrelevant documents [17], compromising precision critical for cell designs.

To address this limitation, we introduce SynBioGPT v2.0, which reduces dependency on vector search and instead harnesses the LLM’s reasoning and analysis capabilities. By decomposing user queries into multiple sub-questions, performing targeted keyword-based searches for each, and synthesizing the retrieved information into a coherent answer, SynBioGPT v2.0 ensures more accurate and contextually relevant responses. This approach minimizes misinterpretation and enhances the platform’s overall reliability.

## Methods

The design of SynBioGPT v2.0 is organized into three distinct stages: data acquisition and cleaning, index creation and storage, and index loading with external service deployment (Figure 1). In the initial phase, open-access PDF documents are systematically gathered, with each document uniquely identified by a standardized DOI and converted into a Hugging Face dataset format for raw data preservation. This raw dataset is then processed using Docling, which converts PDFs into Markdown files while disabling built-in image handling, OCR, and table enhancement options. The resulting Markdown files are incorporated into the dataset. Further refinement is achieved by applying spaCy Layout (https://github.com/explosion/spacy-layout) to extract paragraph structures and remove citations, sensitive information, and headers or footers, thereby generating a curated Markdown column. Additionally, spaCy Layout is employed to extract the table of contents (TOC) from the curated Markdown, storing this information in a separate TOC column. Metadata obtained during the open-access collection process—such as author details, country, abstract, title, and classification—is also expanded and stored as individual columns within the dataset.

**Figure 1.**
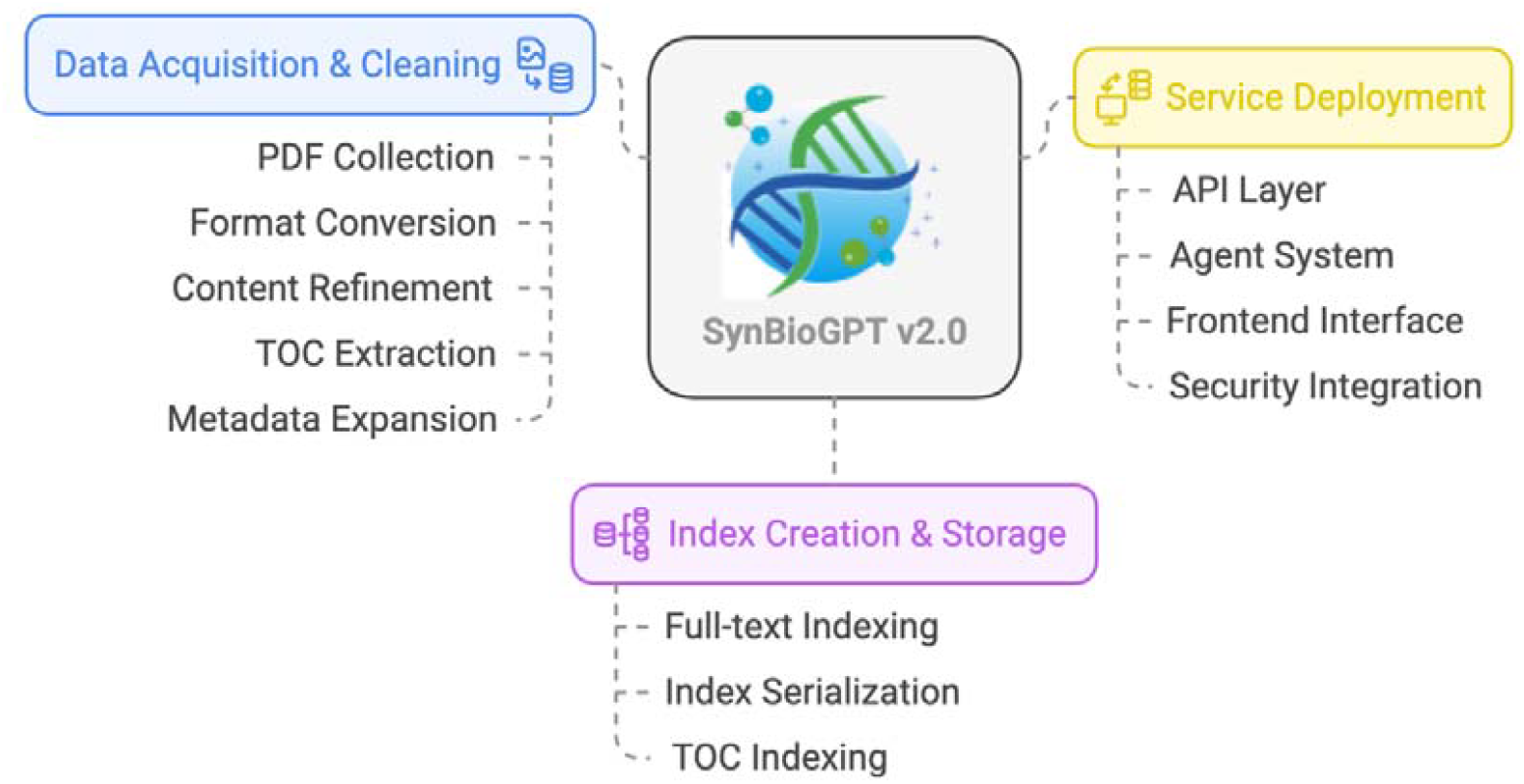
The architecture of SynBioGPT v2.0

In the subsequent phase, SynBioGPT v2.0 establishes its indexing framework. The system computes word frequency data from the curated Markdown using Python’s built-in collections.Counter, and then employs TF/IDF calculations to generate BM25 index data for each article. This index data is serialized locally using Python’s pickle module for efficient storage and future retrieval. A parallel indexing process is performed on the TOC column to create a dedicated directory index file, which is also saved as a pickle file. In the final stage, the platform transitions to providing external services. A FastAPI-based API server is set up to handle user requests, pre-loading both the full-text and TOC index pickle files. Moreover, the initialization of smolagents’ CodeAgent is performed in advance to prevent redundant resource allocation during concurrent requests. For the frontend, Streamlit is used to facilitate user interaction, with session and chat history data stored in a local SQLite database. Finally, integration with the BDC’s single sign-on (SSO) service is implemented through OAuth, ensuring secure third-party account access and robust authentication.

## Results

### Comparative Analysis of SynBioGPT v1.0 and v2.0 in Synthetic Biology Knowledge Retrieval

Both SynBioGPT v1.0 and v2.0 are designed as LLM-based platforms for synthetic biology knowledge retrieval, yet they employ different approaches to balance the trade-offs between comprehensive retrieval and precision. SynBioGPT v1.0 utilizes vector search technology, transforming text into numerical vectors to retrieve documents based on semantic similarity. This method is particularly effective for handling complex or ambiguous queries by uncovering relevant information that might not contain specific keywords. However, its reliance on semantic matching introduces risks, as it may retrieve documents that are semantically similar but contextually irrelevant—an issue that can lead to hallucinations and inaccuracies, especially given the high precision demands of synthetic biology.

In contrast, SynBioGPT v2.0 adopts a more refined information retrieval approach by reducing dependency on potentially error-prone vector search methods. Instead, it decomposes user queries into multiple sub-questions to cover various angles and employs keyword-based searches for each sub-question. This targeted approach involves extracting key paragraphs from the search results, which the LLM then synthesizes into coherent responses. Although keyword-based searches may have a narrower coverage compared to vector search—potentially missing semantically related but non-keyword documents—the method significantly enhances precision and reduces the risk of retrieving irrelevant information. SynBioGPT v2.0 also benefits from an expanded knowledge base that is updated monthly, ensuring continuous access to the latest research and supporting its application in high-precision scenarios such as cell design. Overall, while both versions share the common goal of delivering accurate and reliable synthetic biology knowledge, SynBioGPT v2.0 leverages advanced query decomposition and strategic keyword searches to overcome the limitations observed in SynBioGPT v1.0’s vector search approach.

**Table 1.** Comparative Analysis: v1.0 vs. v2.0.

| Aspect | SynBioGPT v1.0 | SynBioGPT v2.0 |
| --- | --- | --- |
| Retrieval Method | Vector search on pre-built knowledge base | Keyword-based index searches for sub-questions |
| Knowledge Source | Curated 51,777 studies | Updated monthly |
| Hallucination Risk | Higher due to semantic similarity retrieval | Lower, with targeted information retrieval |
| LLM Role | Generates answers from retrieved documents | Generates sub-questions, synthesizes findings |
| Efficiency | Faster with pre-built index | Potentially slower, multiple searches required |
| Transparency | Less explicit, relies on vector similarity | More transparent, sub-questions and sources clear |

### Query Processing and Answer Generation in SynBioGPT v2.0

SynBioGPT V2.0 represents an advancement over its predecessor, designed to capitalize on the enhanced reasoning and analytical capabilities of LLMs. Departing from traditional vector search methodologies, this version employs a keyword-based, sub-question-driven framework to improve both the precision and comprehensiveness of its responses. The core design principles of SynBioGPT v2.0 are outlined as follows:

1. **Utilization of LLM Reasoning Capabilities**: Rather than employing intricate vector search algorithms, SynBioGPT v2.0 harnesses the inherent reasoning strengths of the LLM. This shift simplifies the system architecture while maximizing the model’s ability to interpret and synthesize complex information.
2. **Query Decomposition**: For any user-submitted query, the system generates multiple sub-questions, each addressing a distinct facet of the original inquiry. This decomposition ensures a thorough exploration of the topic, capturing diverse perspectives that might otherwise be overlooked.
3. **Keyword-Driven Search**: Each sub-question is formulated as a keyword-based query and executed against indexed databases or knowledge repositories. This method facilitates precise term matching, enhancing the relevance of retrieved information.
4. **Direct Paragraph Extraction**: Search results yield relevant paragraphs that are extracted in their original form, without additional processing. These unadulterated excerpts are then supplied directly to the LLM, preserving their contextual integrity for subsequent analysis.
5. **Information Synthesis**: The LLM integrates the information gleaned from the extracted paragraphs across all sub-questions, producing a cohesive and comprehensive final answer. This process emulates human research practices, where disparate sources are synthesized into a unified response.

To demonstrate the practical application of this methodology, consider the following example:

### User Query

“How can I optimize the production of L-Lysine in *Corynebacterium glutamicum*?”

The processing workflow of SynBioGPT V2.0 unfolds as follows:

1. **Sub-question Generation**: The LLM constructs a series of sub-questions to address various dimensions of the query:
  - optimizing L-Lysine production in *Corynebacterium glutamicum*
  - metabolic engineering strategies for L-Lysine production in *Corynebacterium glutamicum*
  - genetic engineering for L-Lysine production optimization in *Corynebacterium glutamicum*
  - L-Lysine biosynthetic pathway NADPH supply optimization *Corynebacterium glutamicum*
  - fermentation optimization strategies for L-Lysine production in *Corynebacterium glutamicum*
  - carbon flux optimization reducing by-products L-Lysine production in *Corynebacterium glutamicum*
2. **Keyword Search**: Each sub-question is converted into a keyword query and used to interrogate relevant databases or web resources. For example, a search for “metabolic engineering strategies for L-Lysine production in *Corynebacterium glutamicum*” retrieves articles detailing genetic modifications and metabolic engineering strategies.
3. **Paragraph Extraction**: From the search outputs, key paragraphs are identified and extracted. These may encompass methods sections, results sections, or discussions sections from scientific papers that pertain directly to the sub-questions.
4. **Synthesis by LLM**: The LLM evaluates the collected paragraphs, integrating the findings into a comprehensive response. The resulting answer not only offers actionable insights for optimizing L-Lysine production but also ensures transparency by attributing sources.

This structured approach enables SynBioGPT v2.0 to deliver detailed, well-rounded answers to complex queries, leveraging the LLM’s reasoning prowess to systematically dissect and address multifaceted problems in synthetic biology.

### Performance Evaluation of SynBioGPT v2.0 with Varied Inference Backends and Comparison to v1.0 in Synthetic Biology Tasks

To evaluate the efficacy of LLMs in synthetic biology applications, we employed a test set consisting of 100 questions, including 71 specific, fact-based questions and 29 open-ended, reasoning-based questions. These questions spanned key synthetic biology domains such as gene mutation, overexpression, co-expression, exogenous gene integration, precursor utilization, promoter dynamics, and competitive pathway exploration [16]. In this study, we assessed the performance of SynBioGPT v2.0, a domain-specific LLM, using three distinct inference backends—DeepSeek V3 [18], Gemini-2.0-flash, and Claude-3.7-sonnet—to investigate the impact of backend selection on model outcomes. Furthermore, we benchmarked SynBioGPT v2.0 against its predecessor, SynBioGPT v1.0, which was based on the Llama3-8B-Instruct model and previously established as the top performer among models tested in its iteration [16].

The experimental results demonstrate that incorporating RAG technology significantly enhanced accuracy, achieving an improvement of 26–37% (Figure 2 A). These findings highlight the crucial role of domain-specific context in optimizing model performance. Notably, while the core architecture of SynBioGPT v2.0 remained unchanged across evaluations, its performance on synthetic biology tasks was markedly influenced by the choice of inference backend. Among the tested backends, Claude-3.7-sonnet yielded the highest accuracy, outperforming both DeepSeek V3 and Gemini-2.0-flash (Figure 2 A). This superior performance suggests that Claude-3.7-sonnet possesses enhanced reasoning capabilities and a stronger capacity for integrating domain-specific knowledge, making it particularly adept at addressing the intricate, interdisciplinary nature of synthetic biology queries. In contrast, DeepSeek V3 and Gemini-2.0-flash exhibited elevated error rates, especially on specific questions requiring precise factual recall, indicating potential limitations in their knowledge depth or contextual understanding within this domain.

**Figure 2.**
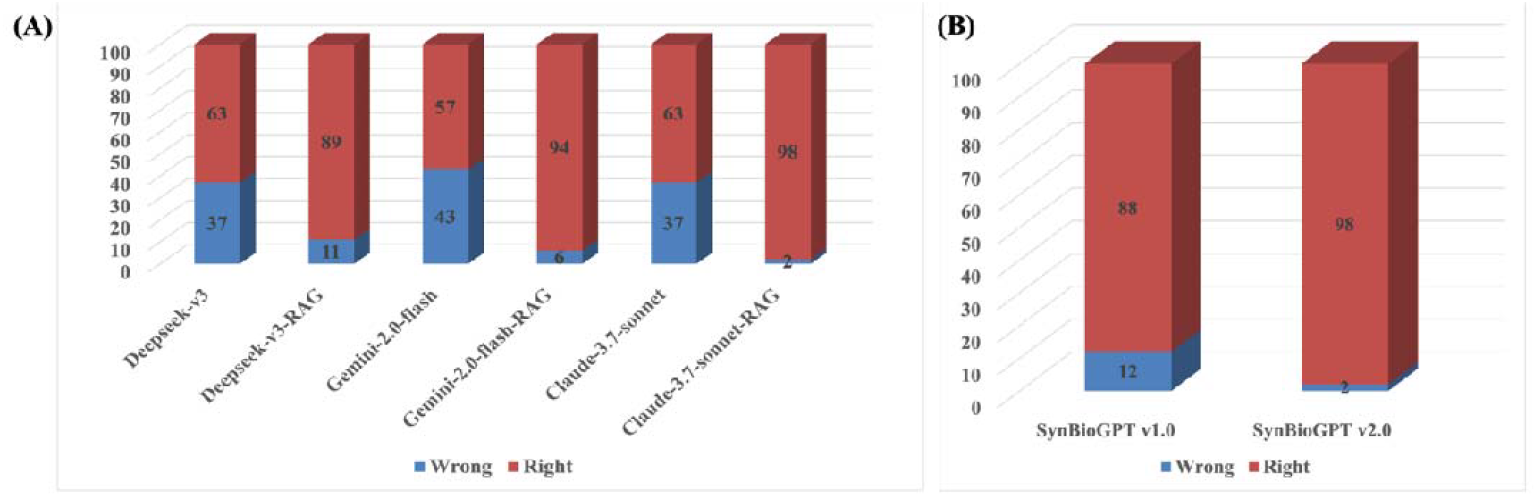
Statistical results of SynBioGPT answering 100 manually curated questions using different models and knowledge bases. (A) Comparison of the performance of DeepSeek V3, Gemini-2.0-flash, and Claude-3.7-sonnet in answering synthetic biology questions with and without retrieval augmentation. (B) The performance comparison between the previous version (SynBioGPT v1.0) and the updated SynBioGPT v2.0.

A comparative analysis of SynBioGPT v2.0 and v1.0 further underscores the advancements in the newer version. Using the same 100-question test set, SynBioGPT v1.0, powered by Llama3-8B-Instruct, achieved an accuracy of 88%, with 12% of responses incorrect [16]. In contrast, SynBioGPT v2.0, when paired with the Claude-3.7-sonnet backend, attained an accuracy of 98%, reducing the error rate to just 2% (Figure 2 B). This 10% improvement highlights the effectiveness of refinements introduced in v2.0, which may include expanded training on synthetic biology-specific datasets, improved handling of complex queries, or optimizations in the model’s interaction with advanced backends. Notably, the 98% accuracy is specific to the Claude-3.7-sonnet configuration, suggesting that the backend plays a critical role in unlocking the full potential of v2.0.

These findings emphasize the interplay between model design and inference infrastructure in achieving high performance in specialized domains. The exceptional results with Claude-3.7-sonnet imply that its underlying architecture or training regimen may better align with the demands of synthetic biology, such as parsing technical terminology or reasoning through multi-step biological processes. Conversely, the relatively weaker performance of DeepSeek V3 and Gemini-2.0-flash on specific questions suggests a need for further investigation into their limitations, potentially related to training data coverage or inference-time reasoning efficiency.

## Discussion

SynBioGPT v2.0 represents a significant advancement over its predecessor through several innovative improvements. Central to these advancements is the introduction of a query decomposition strategy, which dissects complex queries into manageable subproblems. This structured approach, coupled with a shift from vector-based retrieval to a targeted keyword search method, has markedly improved retrieval accuracy. Notably, when using the Claude-3.5-sonnet backend, accuracy increased from 88% in version 1.0 to 98% in v2.0, demonstrating the method’s efficacy in minimizing interference from contextually irrelevant documents.

However, the reliance on keyword search introduces certain limitations. Keyword-based methods depend heavily on exact term matching, which can be problematic in rapidly evolving fields such as synthetic biology. Given the diversity and dynamic nature of the terminology, relevant documents might be overlooked if they do not contain the specific keywords used in the query. For instance, a paper discussing “metabolic flux analysis” may be highly pertinent yet missed if it does not explicitly reference “metabolic engineering.” A hybrid approach that combines keyword search with semantic techniques, such as word embeddings, could significantly improve both precision and recall.

Another critical component of SynBioGPT v2.0 is the generation of subproblems for query decomposition. The effectiveness of this mechanism is heavily contingent on the quality of the generated subqueries. If subproblems are poorly defined, they can lead to retrieval results that deviate from the user’s original intent. It is therefore essential to rigorously evaluate the subproblem generation process. Comparative studies, such as contrasting LLM-generated subproblems with those manually designed or expert-validated, would offer valuable insights.

Finally, the choice of inference backend has a profound impact on overall system performance. Evaluations indicate that the Claude-3.7-sonnet backend outperforms alternatives like DeepSeek V3 and Gemini-2.0-flash, particularly in handling fact-based queries. This performance gap is likely due to differences in model architecture and the diversity of training data. The superior reasoning capabilities of Claude-3.7-sonnet make it particularly well-suited for addressing the complex queries typical in synthetic biology. Future backend selections should continue to prioritize models with strong domain-specific language understanding and reasoning capabilities, with regular benchmarking against emerging models to maintain peak performance.

In conclusion, while SynBioGPT v2.0 has demonstrated substantial improvements in retrieval accuracy and overall system transparency, challenges remain. Addressing the inherent limitations of keyword search, enhancing the evaluation of subproblem generation, ensuring continuous updates of the knowledge base, and optimizing inference backend selection are pivotal for future iterations. Integrating hybrid retrieval approaches and rigorous experimental validations will be key to further enhancing the practical utility and reliability of SynBioGPT in the dynamic field of synthetic biology.

## Conclusion

SynBioGPT v2.0 represents a significant evolution from v1.0, addressing hallucinations and improving accuracy through a structured, keyword-based, sub-question-driven approach. By bridging the adaptability of LLMs with domain-specific knowledges, it establishes a novel paradigm for accelerating DBTL cycles, with potential impacts on both academic research and industrial applications in synthetic biology.

## Author contributions

Z.M., X.L. and H.M. designed the study and wrote the manuscript. J.D, R.W. designed the architecture and deployment of SynBioGPT. H.L. developed the frontend interface of SynBioGPT. Z.M., J.G. and Z.S. collected synthetic biology-related data and conducted testing on SynBioGPT. All authors have read and approved the final manuscript.

## Acknowledgments

This research was supported by the Strategic Priority Research Program of the Chinese Academy of Sciences, the National Natural Science Foundation of China (32300529, 12326611), and Tianjin Synthetic Biotechnology Innovation Capacity Improvement Projects (TSBICIP-PTJJ-012).

